# Modelling the nuclear envelope of *HeLa* cells

**DOI:** 10.1101/344986

**Authors:** Cefa Karabağ, Martin L. Jones, Christopher J. Peddie, Anne E. Weston, Lucy M. Collinson, Constantino Carlos Reyes-Aldasoro

## Abstract

This paper describes a framework for the automatic segmentation of the nuclear envelope of cancerous HeLa cells and the modelling of the volumetric shape against an ellipsoid. The framework is automatic and unsupervised and reported a Jaccard Similarity Index of 0.968 against a manual segmentation. The modelling of the surface provides a visual display of the variations, both smooth and rugged over the surface, and measurements can be extracted with the expectation that they can correlate with the biological characteristics of the cells.

## 1 Introduction

The field of *Computational Pathology* has grown in recent years bringing together computational and mathematical methods that are applied to disease-related data sets [1]. Computational pathology can combine various sources of data such as medical records, laboratory data, genomics, proteomics, and a variety of images with different stainings, antibodies and biomarkers [2, 3]. Some of its aims are to quantify pathological data with various techniques such as machine learning [4] to enable the best possible medical decisions [5]. The automation of the acquisition, especially through the use of whole slide imaging [6] and high-throughput microscopic equipment, has been instrumental in the development field. Labs can now acquire tens of thousands of data sets that can easily exceed gigabytes of data every month [7]. Previous work has applied computer-based image analysis for cell detection and classification [8], tissue classification [9], nuclei and mitosis detection [10], microvessel segmentation [11] and other immunohistochemistry scoring tasks [12] in histopathological images.

Generally, although not exclusively, the images associated with computational pathology are light or fluorescence microscopy and stained through immunohistochemistry. Less frequently considered are images observed with electron microscopy (EM), phase contrast and differential interference contrast (DIC) [13]. One important difference between these last modalities and light and fluorescence, is the fact that the intensity of each pixel or voxel is not exclusively related to the presence or absence of a marker or stain, and thus the elements of interest have intensities above and below that of the background. Another important difference is that EM can provide 3D data sets in a similar way to multiphoton and confocal microscopes, albeit through a destructive process [14, 15]. In addition, in the case of EM, the much higher resolution and magnification changes considerably the problems to solve beyond separating and counting cells and nuclei, or finding their spatial distribution. As there may be a single cell of interest, understanding the nuclear formation [16], the arrangement of chromosomes [17] or the breaching of the nuclear envelope [18] become relevant. The integrity of the nuclear envelope (NE), which separates nucleoplasm and cytoplasm, is of great interest as for some time it has been assumed that the NE brakes down only during mitosis, however, in cases of virus infection or cancer, the NE may remodel outside mitosis [18].

A problem common to all imaging modalities is the determination of a ground truth (GT) against which to validate segmentation results. Despite the significant disadvantages of time and inter- and intra-user variability, manual or semiautomatic delineation of structures like the NE is widely used [19, 20]. Recently, a new approach known as *citizen-science* (CS) [21], has been used to find a GT by leveraging the power of the internet and *an army* of non-experts. The expectation is that out of large enough number of delineations, a useful segmentation with comparable accuracy of that provided by experts can be extracted.

In this paper, a framework to analyse the NE of HeLa cells is presented. The framework extends an automatic segmentation of the NE [22] and models the NE as a 2D surface against a 3D ellipsoid. Distances from the NE to the ellipsoid and the local variation of the distances are calculated with the expectation that they can allow the identification of different biological processes.

## 2 Materials and Methods

### 2.1 Materials: HeLa cells preparation and acquisition

The preparation details of the cell have been published previously, but briefly, wild type HeLa cells were embedded in resin (*Durcupan*) as per the guidelines of the National Centre for Microscopy and Imaging Research (NCMIR) [23].

Images were acquired using a serial blockface scanning electron microscopy (SBF SEM) using a 3View2XP (Gatan, Pleasanton, CA) attached to a Sigma VP SEM (Zeiss, Cambridge). The resolution of the images was 8192 x 8192 pixels over a total of 518 slices, with 10 × 10 × 50 nm voxel size with 0-255 intensity levels. Individual cells were manually cropped as volumes of interest as substacks of 2000 × 2000 × 300 voxels (Figs. 1a,d) and were saved as single channel *tif* files.

**Fig. 1:**
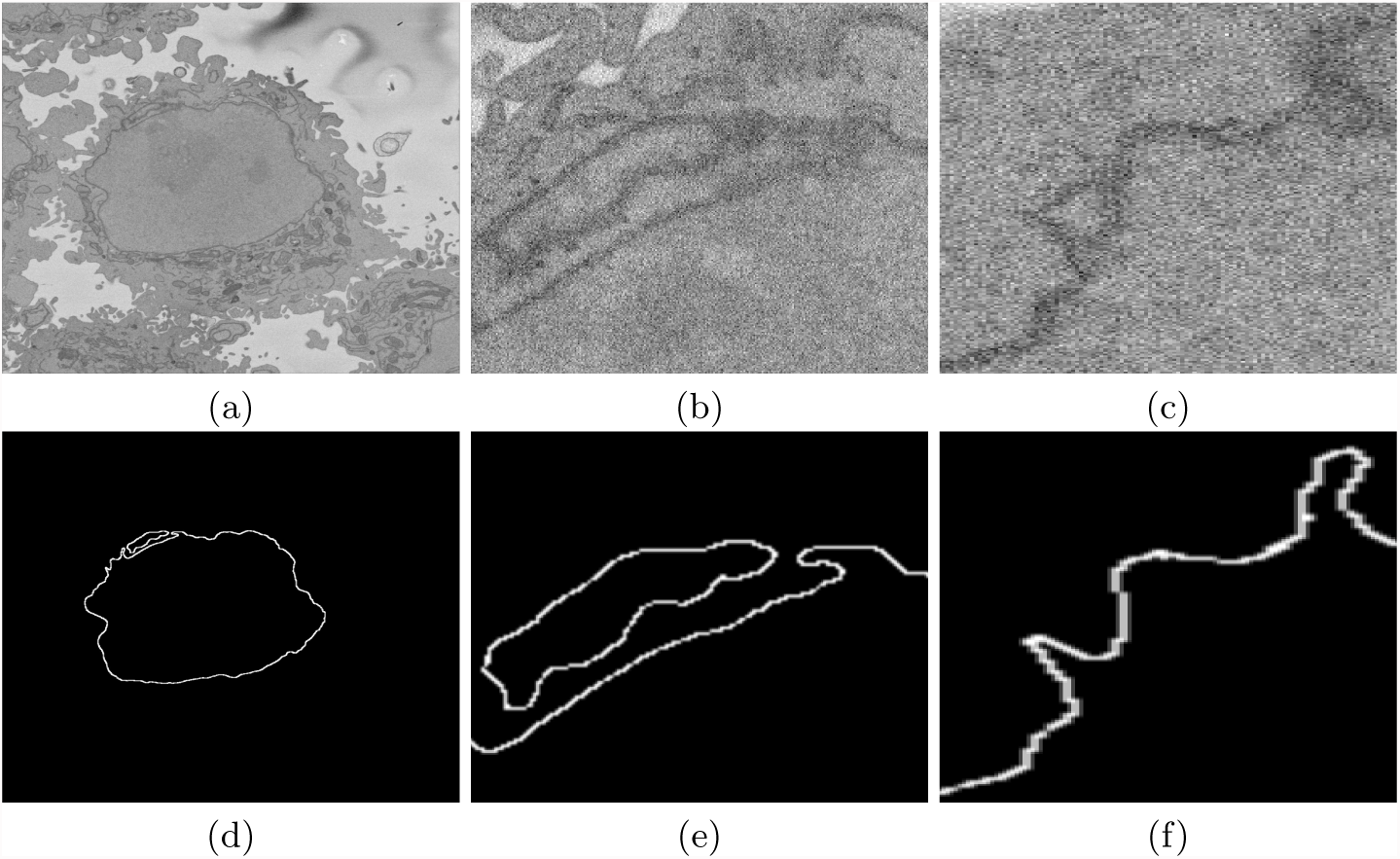
(a) One representative EM image containing one HeLa cell at the centre and fragments of other cells surrounding it. (b,c) Two regions of interest (ROI) which illustrate the difficulty of identifying the nuclear envelope. (d) Manual delineation of the nuclear envelope performed by a single expert. (e,f) Delineations of the ROIs from (b,c). Notice in (b,e) the disjoint region that the expert considered to belong to the nucleus and in (c,f) the uncertainty in some regions of the boundary.

### 2.2 Ground Truth

Manual delineation of every slice of the data set was performed by an expert who had no further input in the processing of the data. As the HeLa cells can have complicated 3D shapes, in some cases it was necessary for the expert to scroll up and down the slices to determine if a certain region, which appeared disjoint in a certain slice, was part of the nucleus (Figs. 1b,e).

### 2.3 Automatic segmentation of the nuclear envelope

The NE was segmented following the framework described in [22], which is available open-source as Matlab code (https://github.com/reyesaldasoro/HeLa-CellSegmentation). The framework exploited the darker intensity of the NE as compared with the cytoplasm and the nucleoplasm by Canny edge detection. The edges were then dilated to connect disjoint edges, which were part of the NE, due to intensity variations of the envelope itself (Fig. 2a). The connected pixels not covered by the dilated edges were labelled to create a series of superpixels (Fig. 2b). The superpixel size was not restricted as large superpixels covered the areas of background. Morphological operators were used to: remove regions in contact with the edges of the image, remove small regions and fill holes inside larger regions (Fig. 2c). The central superpixel was selected as the nucleus and further morphological operators were applied to close the jagged edge. Sensitivity analyses to determine the optimal parameters were performed (results not shown).

**Fig. 2:**
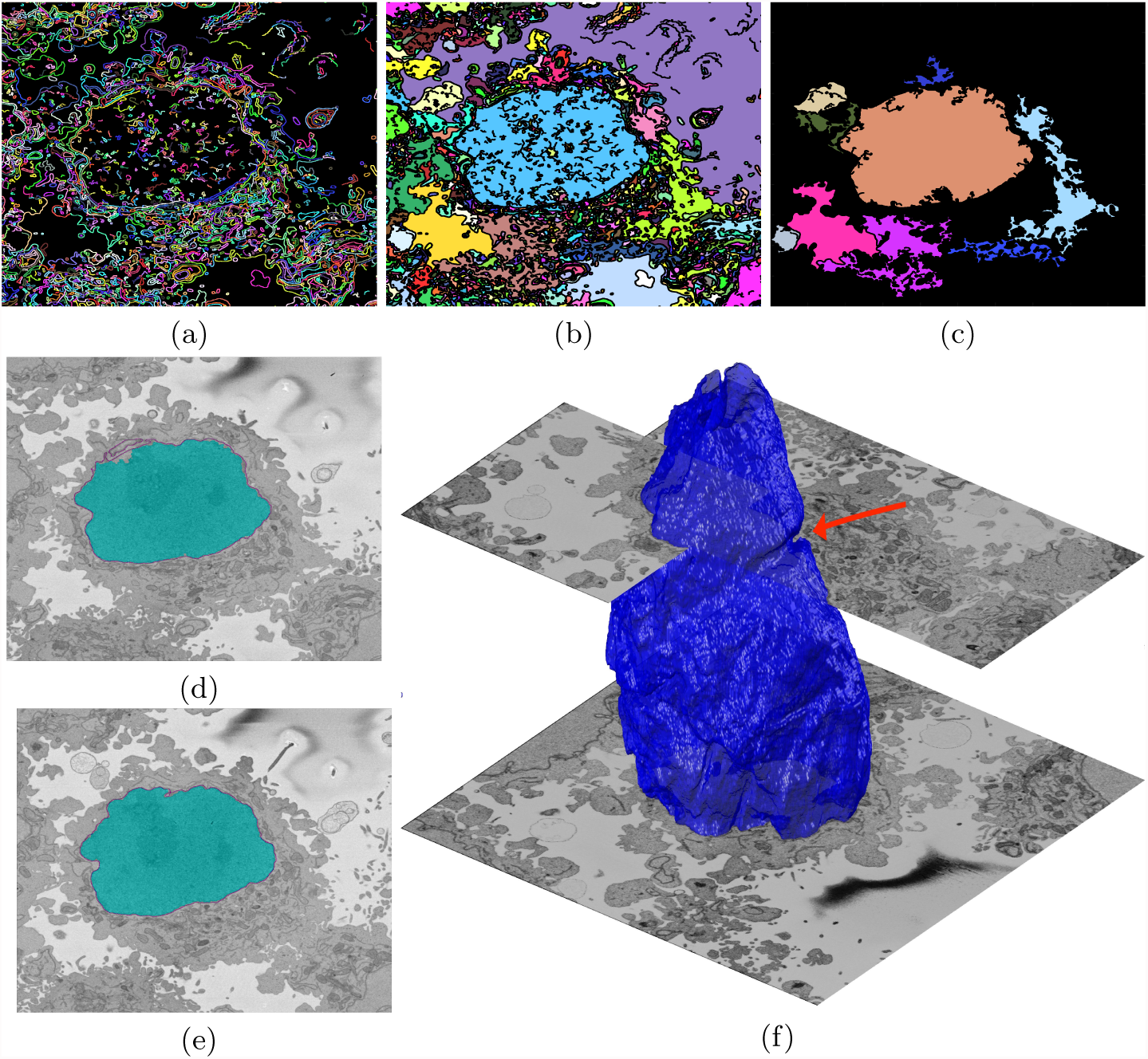
Illustration of the pipeline to segment the nuclear envelope of the HeLa Cell. (a) Edges detected by Canny algorithm. (b) Labelled superpixels. (c) Morphological processing and selection of large superpixels. The edges and the labelled areas have been assigned random colours for visualisation purposes. (d,e) Segmentation of two slices (cyan shades) and the manual segmentation (magenta line). (f) Surface of the NE and partial EM slices to give context. Notice the notch (arrow) on the upper right side of the NE.

### 2.4 Validation of the segmentation

In order to compare the results of the automated segmentation with the manually segmented GT, the Jaccard Similarity Index (JI) [24] of intersection over union of areas was calculated. For each slice, the manual delineation was morphologically closed to generate a region rather than a line.

### 2.5 Nuclear envelope shape modelling

To further study the shape of the segmented NE, this was modelled against a 3D ellipsoid. The ellipsoid was adjusted to have the same volume as the nucleus. The surfaces of the ellipsoid and the nucleus were subsequently compared by tracing rays from the centre of the ellipsoid and the distance between the surfaces for each ray was calculated. It was assumed that when the nucleus surface was further away from the centre, the difference was positive. Processing and visualisation were performed in MATLAB^®^ (The Mathworks™, Natick, USA).

### 3 Results

The automatic algorithm produced a rather accurate segmentation of the NE. The JI of the automatic results against the manual GT was 0.968. This value indicate a rather high overlap considering that the shape of the nucleus is very irregular on the upper and lower slices. Furthermore, the shape of the manual delineations can give rise to ambiguities in the assessment of the segmentation correctness. In particular, the delineation connected in different points when invaginations of the NE were present, and thus the GT, which was morphologically closed, was itself incorrectly estimated (Fig 3).

**Fig. 3:**
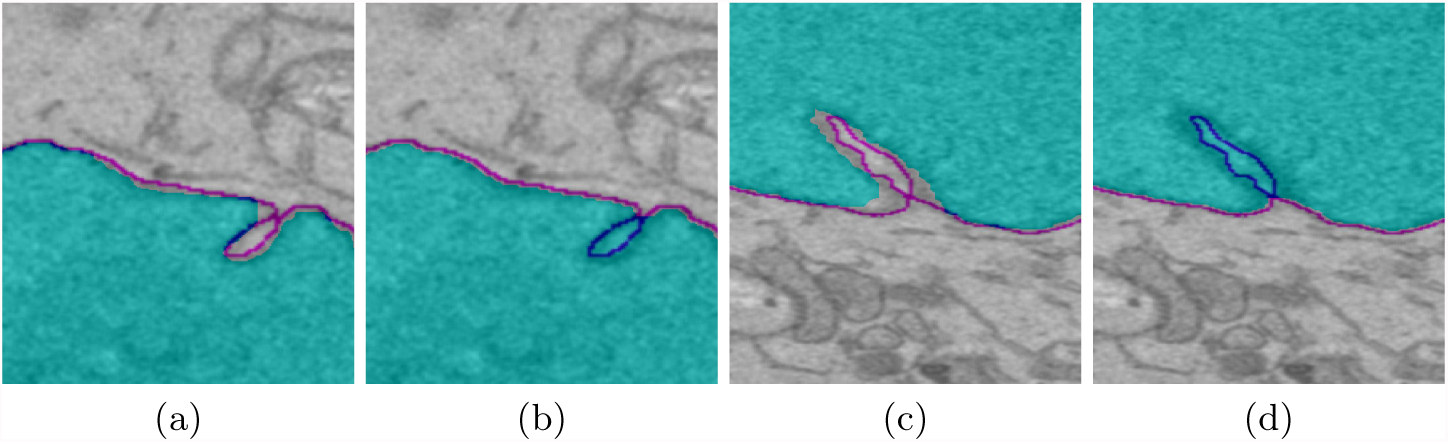
Assessment of the automatic segmentation and the ground truths of two ROIs. In all cases the magenta line corresponds to the manual delineation and the shaded region in blue corresponds to: (a,c) Automatic segmentation, (b,d) Manual GT. Notice how the GT has been incorrectly estimated in the cases where an invagination of the boundary creates a closed loop.

The comparison between the model ellipsoid and the NE reported a JI of 0.7184 (Fig. 4a). This value could be used to assess the roundness or not of a nucleus, and it is speculated that the JI could be related with biological characteristics of cells. This will be validated in the future against a large number of cells. In addition, the measurements of distance from the nucleus to the ellipsoid showed rougher and smoother regions (Figs. 4b,c, 5).

**Fig. 4:**
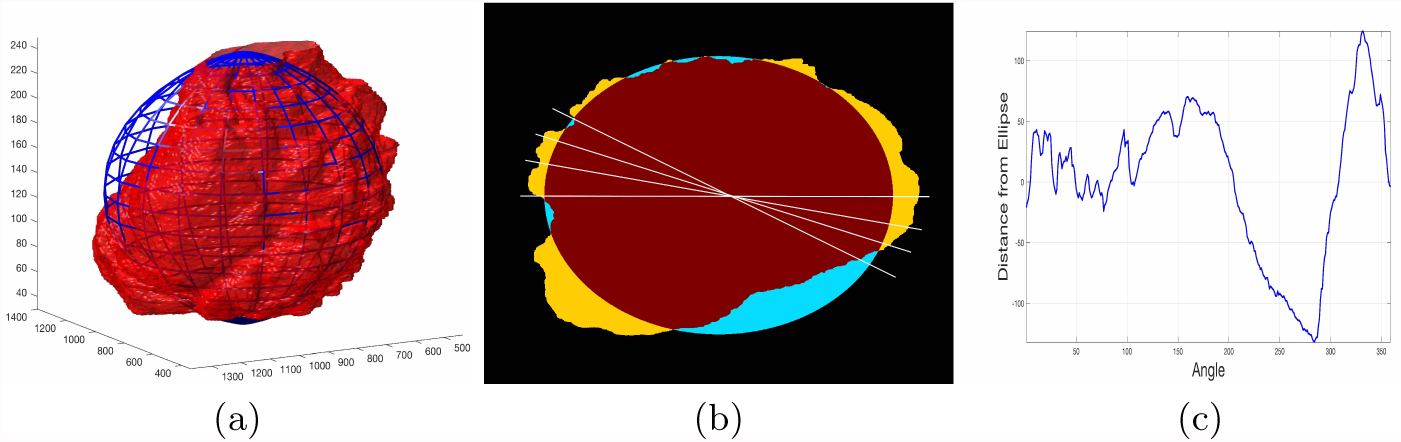
(a) Rendering of the nuclear envelope (red surface) against the model ellipsoid (blue mesh). (b) Illustration of distance calculations by ray tracing in one slice. Yellow regions correspond to the nucleus outside the ellipsoid, cyan regions where nucleus inside the ellipsoid. (c) Measurements obtained along the boundary.

**Fig. 5:**
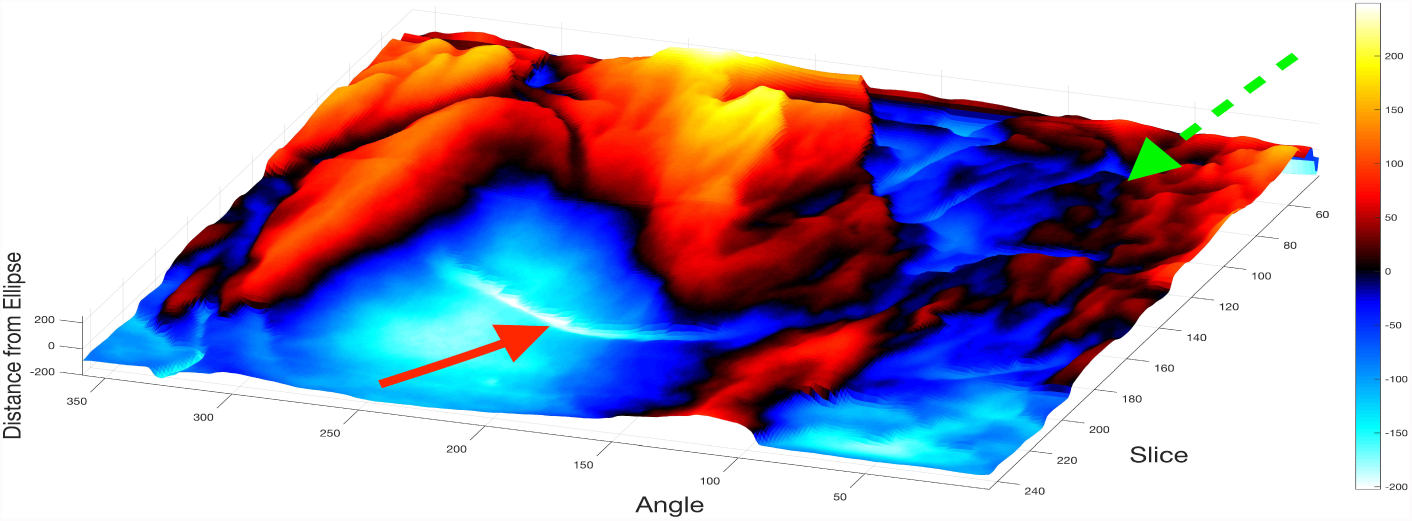
Surface corresponding to the distance from the nuclear envelope to a model ellipsoid. Solid red arrow indicates a notch, dashed green arrow shows rugged region.

The surface corresponding to the distance from the nuclear envelope to a model ellipsoid (Fig. 5) showed graphically the hollow and prominent regions of the cell, but more important, elements such as a notch (solid red arrow) or ruggedness (dashed green arrow) can be an indication of NE brake down or remodelling. The local variation of the surface (Fig. 6) enhanced these elements as the trends were removed. Notice again the notch and the high frequency variation of the rugged region.

**Fig. 6:**
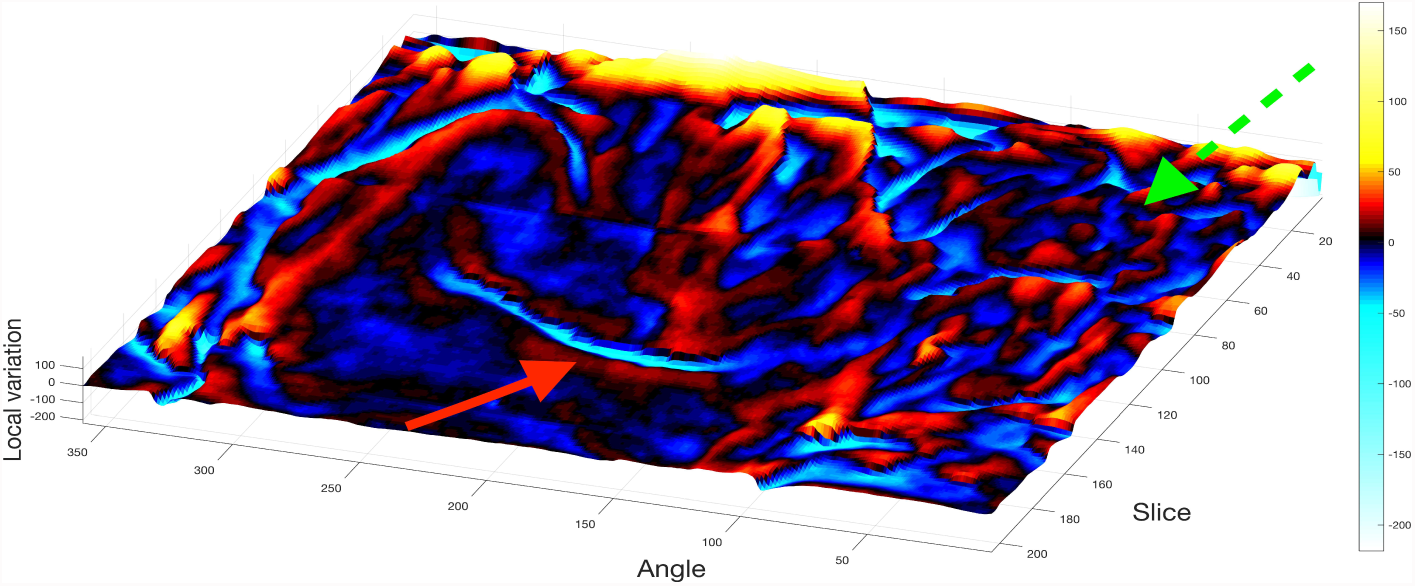
Surface of the local variation of the distances. In both cases, hot shades correspond to positive values and cool shades to negatives. Solid red arrow indicates a notch, dashed green arrow shows rugged region.

## 4 Discussion

The modelling of the NE surface against an ellipsoid can reveal interesting characteristics of the nucleus and the nuclear envelope of a HeLa cancer cell. The 2D maps of the NE surface provides an easier way to assess the characteristics of a 3D structure. Whilst further experimentation with more data sets, and a careful processing of the GTs, is necessary, the framework here described can be useful, as it is fully automated, unsupervised, segmented each slice in approximately 8 seconds and does not require training data.

Future work will concentrate in the segmentation and modelling of a large number of cells, not all of which have a GT segmented by an expert and will be compared against results gathered from the CS approach. For this purpose, a CS project called *Etch a Cell* [25] was created to gather manual segmentations of the NE of HeLa cells.

## 5. Acknowledgements

This work was supported by the *Francis Crick Institute* which receives its core funding from Cancer Research UK (FC001999), the UK Medical Research Council (FC001999), and the Wellcome Trust (FC001999). The authors acknowledge the *Alan Turing Institute* Data Study Groups and Dr Sebastian Vollmer.

